# Amino acid substitutions in human growth hormone affect coiled-coil content and receptor binding

**DOI:** 10.1101/2021.12.16.473085

**Authors:** Andrei Rajkovic, Sandesh Kanchugal, Eldar Abdurakhmanov, Rebecca Howard, Astrid Gräslund, Sebastian Wärmländer, Joseph Erwin, Helena Danielson, Samuel Flores

**Affiliations:** Department of Cell and Molecular Biology; Chemistry, Uppsala University; Department of Biochemistry and Biophysics, Stockholm University; Department of Animal Breeding and Genetics, Swedish University of Agricultural Sciences

## Abstract

The interaction between human Growth Hormone (hGH) and hGH Receptor (hGHR) has great relevance to human diseases such as acromegaly and cancer. HGH has been extensively engineered by other workers to improve binding and other properties. We used a computational screen to select substitutions at single hGH positions within the hGHR-binding site. We find that, while many successfully slow down dissociation of the hGH-hGHR complex once bound, they also slow down the association of hGH to hGHR. We are particularly interested in E174 which belongs to the hGH zinc-binding triad, and which spans coiled-coil helices and obeys the coiled-coil heptad pattern. Surprisingly, substituting E174 with A leads to substantial increase in an experimental measure of coiled-coil content. E174A is known to increase affinity of hGH against hGHR; here we show that this is simply because the off-rate is slowed down more than the on-rate, in line with what has been found for other affinity-improving mutations. For E174Y (and mutations at other sites) the slowdown in on-rate was greater, leading to decreased affinity. The results point to a link between coiled-coiling, zinc binding, and hGHR-binding affinity in hGH, and also suggest rules for choosing affinity-increasing substitutions.

## Introduction

Human Growth Hormone (hGH) binds a single hGH Receptor (hGHR) using its Site 1, a large, physicochemically diverse binding region. It then recruits a second hGHR (Figure 1) to bind at its lower-affinity Site 2. (Cunningham *et al*., 1991b) The hGHR dimerization initiates signaling through the JAK/STAT pathway. Thus one strategy to disrupt signaling is to prevent dimerization. This in turn can be done by destroying binding at site 2, which is easily effected with mutations such as G120K. (Flyvbjerg *et al*., 1999) It is also useful to simultaneously strengthen binding at site 1. (Lowman and Wells, 1993) Pegvisomant, an anti-acromegaly therapeutic, is an hGH variant which was affinity matured at site 1 using phage display, reaching an affinity 400-fold higher than wild type (Lowman and Wells, 1993) However, this generated substitutions at 15 amino acid positions in site 1, and other considerations required manually selected reversions and mutagenesis. The G120K mutation was also generated. Lastly, the protein was PEGylated to extend serum half-life. These manipulations resulted in significant reduction of affinity compared to wild type.(Ross *et al*., 2001) As a result, there remains significant potential for hGH variants to be used as therapeutics and diagnostics for several cancers and growth disorders, hence motivation to understand the biophysics of binding at site 1. We are interested in the possibility of choosing amino acid substitutions at a small number of residue positions, obtaining higher affinity while also working within constraints such as maintaining solubility, enabling modifications such as PEGylation, and avoiding patented substitutions.

**Figure 1.**
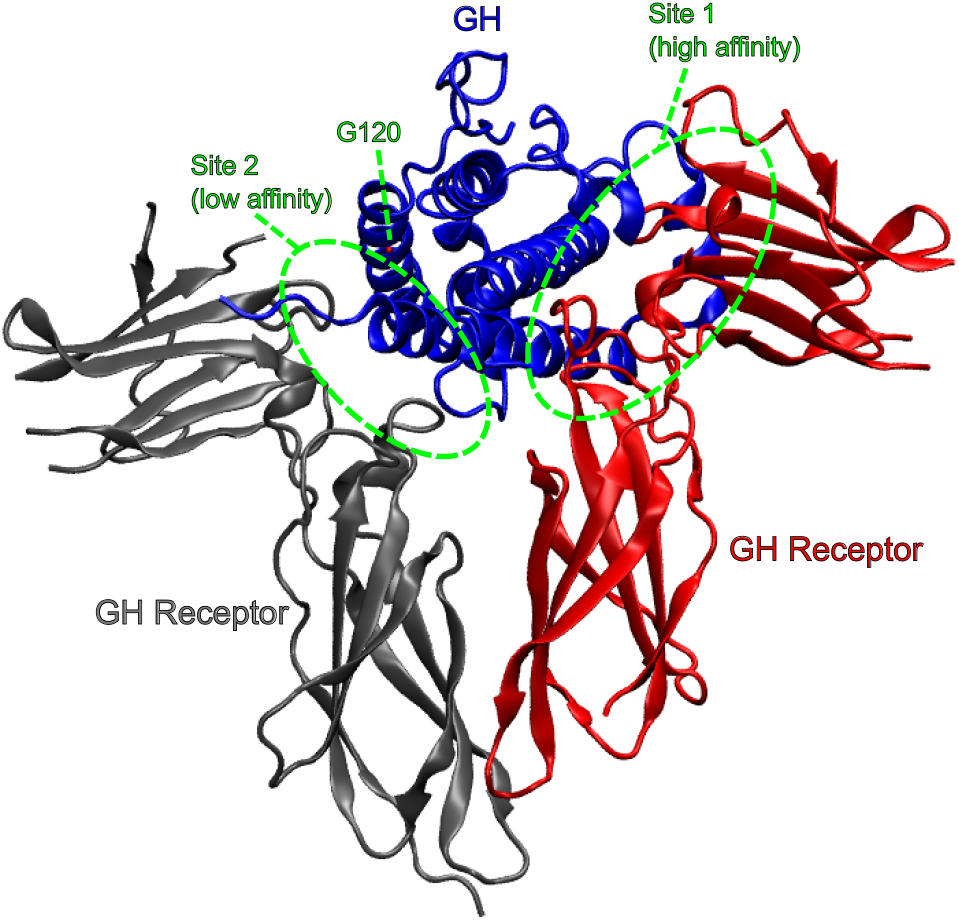
Growth Hormone (GH) initiates signaling by first binding one GH Receptor at its high-affinity Site 1, then recruits a second GHR at its low-affinity Site 2. A GHR antagonist can be made by destroying GHR-binding affinity at Site 2, e.g. with mutation G120K, thus preventing the GH from dimerizing GHR. It is also desirable for Site 1 binding affinity to be increased, to better compete against native GH.

We are also interested in a remarkable result of early work leading up to Pegvisomant development. Alanine scanning is a technique used to map binding interfaces in the absence of structural data. Alanine is smaller than all other canonical amino acids except glycine. Therefore protein-protein interactions (PPIs) are usually weakened if the substitution is at the interface. However E174A is an exception – it increases affinity.(Cunningham and Wells, 1989) E174, along with H18 and H21, is part of a zinc-binding triad (Cunningham *et al*., 1991a) (Figure 2). E174 is also packed between two helices in a coiled-coil and follows the typical heptad repeat. (Lupas *et al*., 1991) We therefore paid particular attention to amino acid E174. How could an amino acid which is part of the zinc binding triad and is packed in a coiled-coil interface, be mutated (to alanine, no less) and increase affinity?

**Figure 2.**
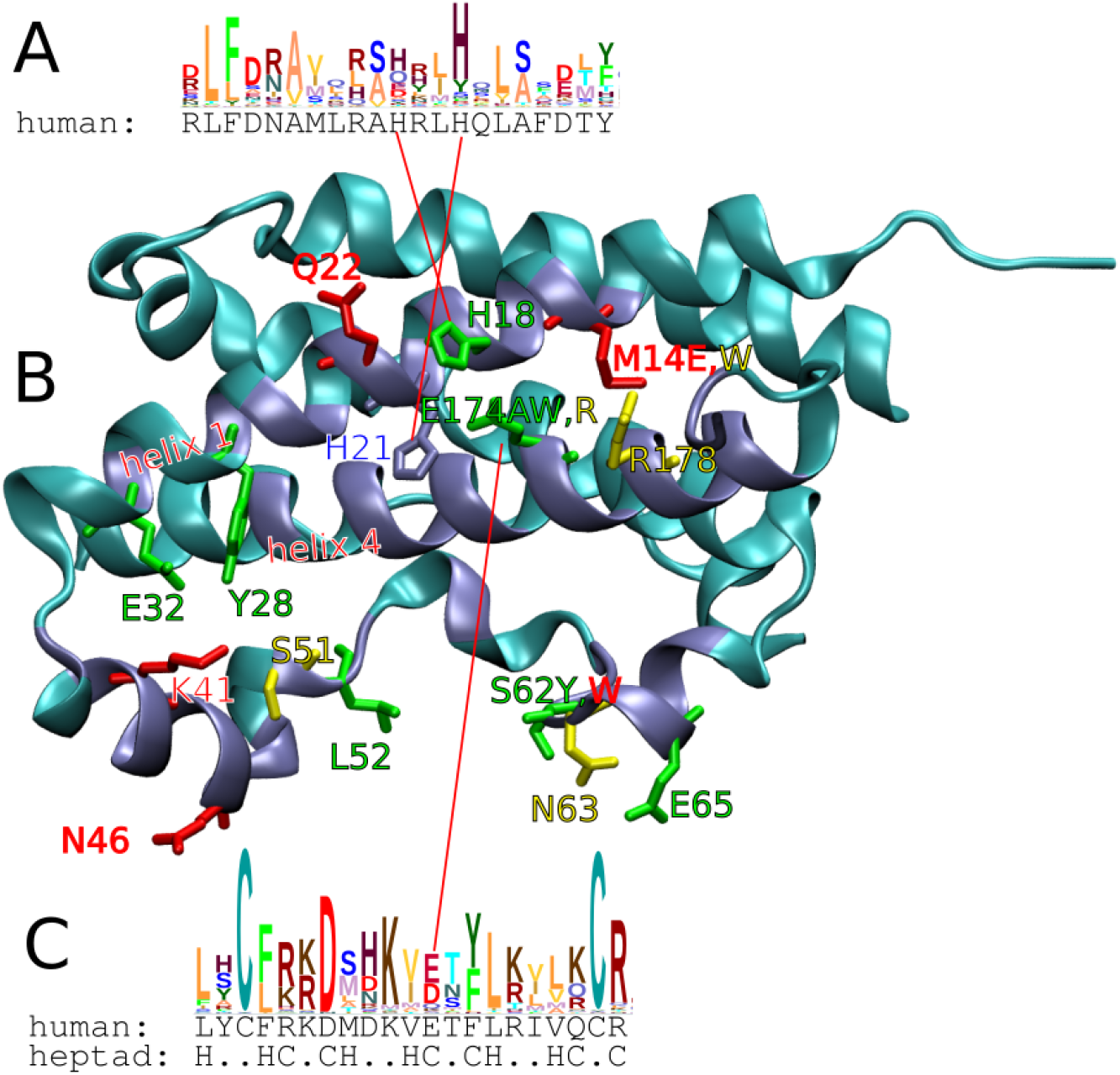
Conserved position E174 is part of the zinc binding triad and obeys the heptad pattern. Coiled-coils tend to follow the heptad pattern of HxxHCxC (H = hydrophobic, C = charged). (A) Helix 1 appears coiled with its antiparrallel partner helix 4. However the former does not follow the heptad pattern, and no coiled-coil signal was found by COILS, MultiCoil, or other expasy coiled-coil predictors. H18 and H21 are conserved (profile generated with Skylign, vertical size proportional to information) and part of the zinc binding triad. Thus the triad appears important for coiled-coiling (B) in the hGH monomer. Site 1 residues are shown in iceblue, remaining residues in cyan cartoons. At several positions, we generated mutations with binding kinetics similar to those of WT (green labels). We identified a second cluster of mutations with similar koff, but much slower kon (red labels). Mutants with kinetics that did not fall into either cluster are labelled in yellow. At some positions different substitutions were tested which produced different kinetics (M14E,W, S62Y,W, E174A,W,R). (C) Helix 4 follows the heptad pattern closely, and is predicted to be in a coiled-coil by COILS. (Lupas et al., 1991) E174 is conserved and part of the zinc binding triad.

Lastly, we are interested in the general problem of computationally engineering proteins, especially to improve binding affinity (Flores *et al*., 2021). Computational-experimental methods have yielded considerable improvements in affinity, but only because they included an experimental high-throughput screening and affinity maturation stage (Whitehead *et al*., 2013). The required expenditure is out of reach of many labs. The end product typically has substitutions at multiple positions, as occurred in (Lowman and Wells, 1993), making it difficult to know which substitutions were most important and why. This lack of guiding knowledge is the likely reason some modifications of (Ross *et al*., 2001) were detrimental to binding. If one could start with a given scaffold, generate a small number of substitutions at single positions, and obtain improvements in affinity for even a small number of these, this would mean a better ability to 1) fine-tune protein properties without immunological, off-target and other unintended effects, 2) avoid patent-protected modifications, and 3) understand the evolutionary purpose of the native scaffold sequence.

We computationally selected 21 substitutions at several single-amino-acid positions, including position 174, expressed the corresponding hGH variants, and measured their effect on hGH-hGHR binding. The results have interesting consequences for protein engineering in general, and for the biophysics of position E174 in particular.

## Method

We used FoldX to identify single-residue substitutions in hGH which have potential to increase affinity to hGHR. We expressed and purified a number of such variants, and measured their hGHR binding kinetics by Surface Plasmon Resonance (SPR). Lastly, we measured the Circular Dichroism (CD) spectra for WT, E174A, E174Y, and L52F.

### Computational selection of mutations

We selected single-residue substitutions most likely to increase affinity, by computational saturation mutagenesis over site 1. Site 1 was defined as all hGH residues within 6.0Å of hGHR, in the Protein Data Bank (PDB) coordinate set 1A22. (Clackson *et al*., 1998) We computed ΔΔG_predicted_ for all 19 possible aa substitutions at all site 1 positions. This was done using FoldX (Guerois *et al*., 2002) and 1A22.(Dourado and Flores, 2014) WT and mutant structures were processed using the RepairPDB, BuildModel, and AnalyzeComplex commands. The results were ranked by ΔΔG_predicted_. Where more than one substitution at a given aa position had a low ΔΔG_predicted_, we kept at most one substitution from each of the following groups: hydrophobic, positively charged, negatively charged, and uncharged polar. Thus at most four substitutions were tested at any residue position.

### Expression of hGH variants in E. coli

The selected hGH variant DNA was generated by gene synthesis and cloned into pET-28b vectors by Genscript. The sequence included a hexahistidine tag at the N-terminus, in order to facilitate purification using a nickel column. Our “WT” sequence was thus:

HHHHHHFPTIPLSRLFDNAMLRAHRLHQLAFDTYQEFEEAYIPKEQKYSFLQNPQTSLC FSESIPTPSNREETQQKSNLELLRISLLLIQSWLEPVQFLRSVFANSLVYGASDSNVYD LLKDLEEKIQTLMGRLEDGSPRTGQIFKQTYSKFDDALLKNYGLLYCFRKDMDKV**E**TFL RIVQCRSVEGSCG Where for reference, the “E” in bold type is E174.

In all cases, recombinant protein expression was carried out in XJB BL21(DE3) cells. N-terminal His6–hGH was expressed by growing cells in LB until mid-exponential phase and induced with 1 mM IPTG (isopropyl-β-D-thiogalactopyranoside). Protein expression was completed with overnight growth at 16°C for solubility, per (Kim *et al*., 2013). Cells were harvested after two rounds of pelleting at 4,500 × *g* for 10 min. Lysis of cell pellets and all subsequent purification were carried out at room temperature, unless otherwise noted. Cell pellets were resuspended in 10 mL of lysis buffer (50 mM Tris-HCl [pH 8.0], 100 mM NaCl, Roche Complete protease inhibitor) and lysed by a cell disruptor (Constant Systems Ltd, UK). Lysate clarification was achieved after 45 mins of centrifugation at 16,000 rpm with temperature set to 4°C. Clarified lysate was filtered through a 0.45 µm membrane. Prior to loading the clarified lysate, the gravity column with 1ml of Ni-Sepharose resin was washed with 10 column volumes of ddH_2_O and equilibrated with 10 column volumes of lysis buffer (10 mM Tris-HCl [pH 7.4], 500 mM NaCl, 5 mM imidazole). Lysate was applied directly to the capped gravity column and set on a shaker at 4°C to gently mix for 40 minutes. After 40 minutes, lysate was eluted and a series of wash steps followed. The column was first washed with 5 column volumes of lysis buffer supplemented with 40 mM imidazole and then washed with 5 column volumes of lysis buffer supplemented with 10mM imidazole. The stepwise gradient was repeated a total of two times. N-terminal His6-hGH was eluted into 1 mL fractions using 5 column volumes of 250 mM imidazole in lysis buffer. Protein purity was assessed by Commassie staining and fractions with the lowest impurities were pooled and concentrated to 250 µL using Amicon Ultra 4 mL Centrifugal Filters. A second round of purification was carried out via size exclusion chromatography, using the Agilent infinity 1220 HPLC system. The fractions containing protein were evaluated for purity using SDS gel electrophoresis and Coomassie staining. All fractions containing only hGH were then pooled together and dialyzed at 4°C in 10 mM HEPES, 150 mM NaCl, 3 mM EDTA, 0.005% Tween 20, pH 7.4. After 24hrs of dialysis, protein was concentrated to 50 µL through the Amicon® Ultra 0.5 mL Centrifugal Filters. The concentration of final growth hormone purified using this method was assessed using a Bradford assay. The purified growth hormone was then flash-frozen for storage.

### Interaction analysis using surface plasmon resonance (SPR) biosensor

The experiments were performed using a Biacore 3000 instrument (Cytiva, Uppsala, Sweden) at 25C. The immobilization of hGHR was carried out by a standard amine coupling procedure on a CM5 biosensor chip (Cytiva, Uppsala, Sweden). hGHR was diluted to 100 µg/ml in sodium acetate buffer, pH 4.5. The CM5 chip surface was activated by an injection of a 1:1 mixture of EDC and NHS for 7 min, at a flow rate of 10 µl/min. hGHR was injected over the activated surface at a flow rate of 2 µl/min until reaching an immobilization level of 1400 - 2700 RU. Then, the surface was deactivated by the injection of 1 Methanolamine for 7 min. After immobilization, a concentration series of hGH, ranging from 0.3 to 10 nM, was injected over the surface, at a flow rate 30 µl/min. An association phase was monitored for 60 sec and a dissociation phase for 240 sec. The surface was regenerated after each cycle by injecting 4.5 M MgCl2 for 2 min. The data was analyzed using Biaevaluation Software, v. 4.1 (Cytiva, Uppsala, Sweden). Sensorgrams were double-referenced by subtracting the signals from a reference surface and the average signals from two blank injections and fitted to a 1:1 Langmuir binding model (Danielson, 2009).

### Circular Dichroism (CD) measurements

CD spectroscopy was used in order to identify the impact of each mutation on the coiled-coiled domain of hGH. Purified protein samples were dialyzed against 2 L of 20 mM sodium phosphate buffer at pH 7.2 overnight in order to remove the tris buffer used during purification. This dialyzed sample was dialyzed a second time overnight against 2 L of fresh 20 mM sodium phosphate buffer at pH 7.2 to completely remove tris buffer.

The dialyzed sample was then loaded into a 10 mm length quartz cuvette. The sample was then loaded into the spectrophotometer (Chirascan). CD spectra between 240-190 nm were collected every two degrees centigrade as the sample was heated from 20-90 C, and then cooled back down to 20 C. The ratio of the measured ellipticity at 222 nm to the ellipticity at 208 nm was used to measure the coiled-coiled nature of each mutant tested in this manner, as well as the wildtype protein, and its change over varying temperatures could be used to approximate the stability of the coiled-coil domain in each variant of hGH.

## Results

### FoldX calculations

The FoldX calculations are summarized in Figure 3. Multiscanning plot, showing ΔΔG_predicted_ using FoldX on the basis of PDB coordinate set 1A22. WT residue type (in single letter code) and residue number is along the bottom axis, for all site 1 residues. Labels along the left axis designate substituted residue type. Dashed horizontal lines represent 0 and –1.0 kcal/mol for the substituted residue type. Dots represent ΔΔG_predicted_ for the indicated residue position and substituted residue type. ΔΔG_predicted_ between –0.5 and –1.0 kcal/mol, are surrounded with a square. ΔΔG_predicted_ < -1.0 kcal/mol, are surrounded with a circle. Thus circled substitutions have the highest priority for experimental ΔΔG measurement, followed by squared substitutions. Note that some positions, e.g. E65, have many low-ΔΔG_predicted_ substitutions, and it is not sensible to test all, so at most we tested the best one of each physicochemical class : hydrophobic, polar uncharged, positively charged (red shading), and negatively charged (blue shading).. At some positions, e.g. E65 and E174, there were many substitutions with low ΔΔG_predicted_ (see circled points), and in those cases we selected the lowest ΔΔG_predicted_ for each of the four physicochemical classes.

**Figure 3.**
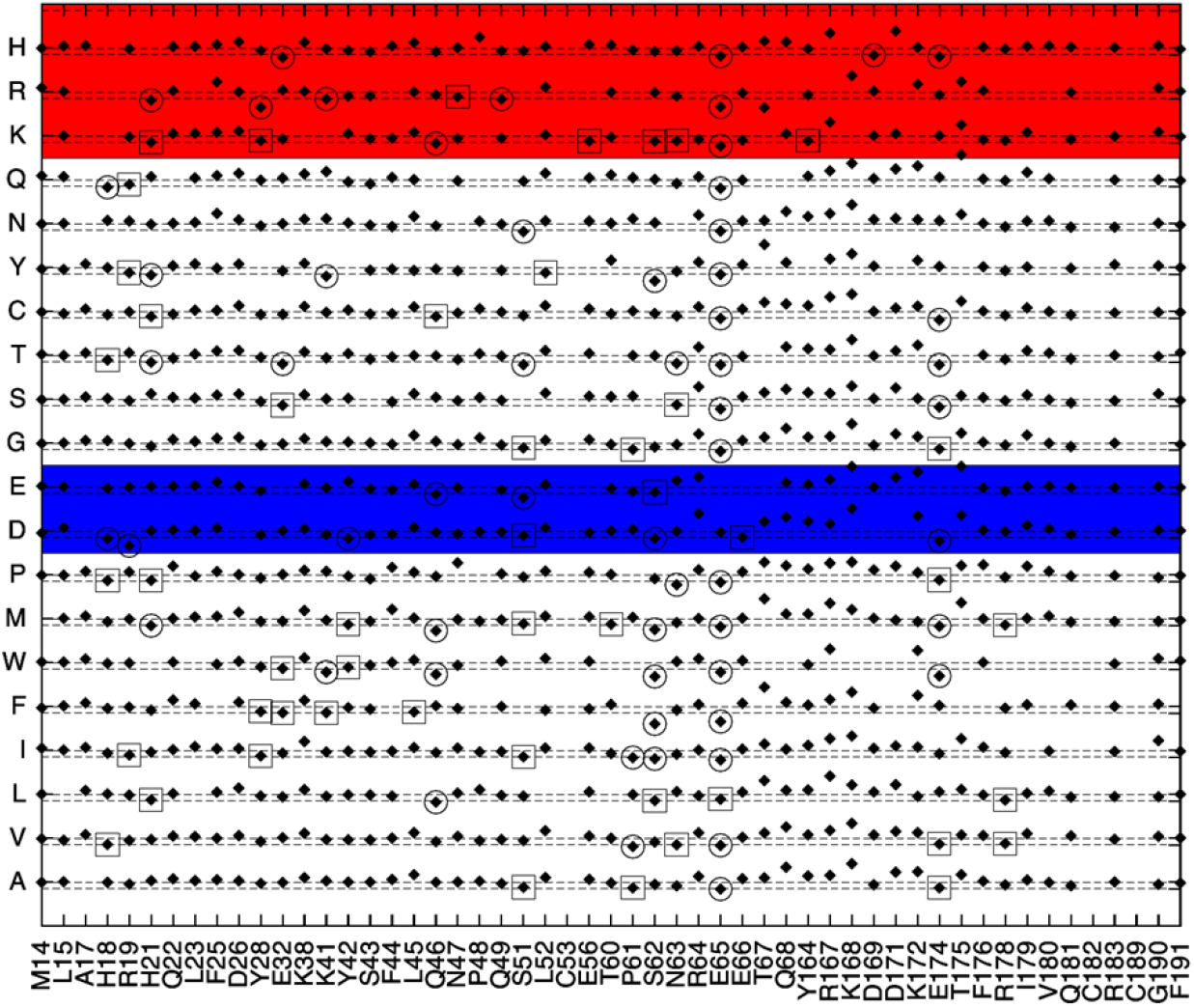
Multiscanning plot, showing ΔΔG_predicted_ using FoldX on the basis of PDB coordinate set 1A22. WT residue type (in single letter code) and residue number is along the bottom axis, for all site 1 residues. Labels along the left axis designate substituted residue type. Dashed horizontal lines represent 0 and –1.0 kcal/mol for the substituted residue type. Dots represent ΔΔG_predicted_ for the indicated residue position and substituted residue type. ΔΔG_predicted_ between –0.5 and –1.0 kcal/mol, are surrounded with a square. ΔΔG_predicted_ < -1.0 kcal/mol, are surrounded with a circle. Thus circled substitutions have the highest priority for experimental ΔΔG measurement, followed by squared substitutions. Note that some positions, e.g. E65, have many low-ΔΔG_predicted_ substitutions, and it is not sensible to test all, so at most we tested the best one of each physicochemical class : hydrophobic, polar uncharged, positively charged (red shading), and negatively charged (blue shading).

**Figure 4.**
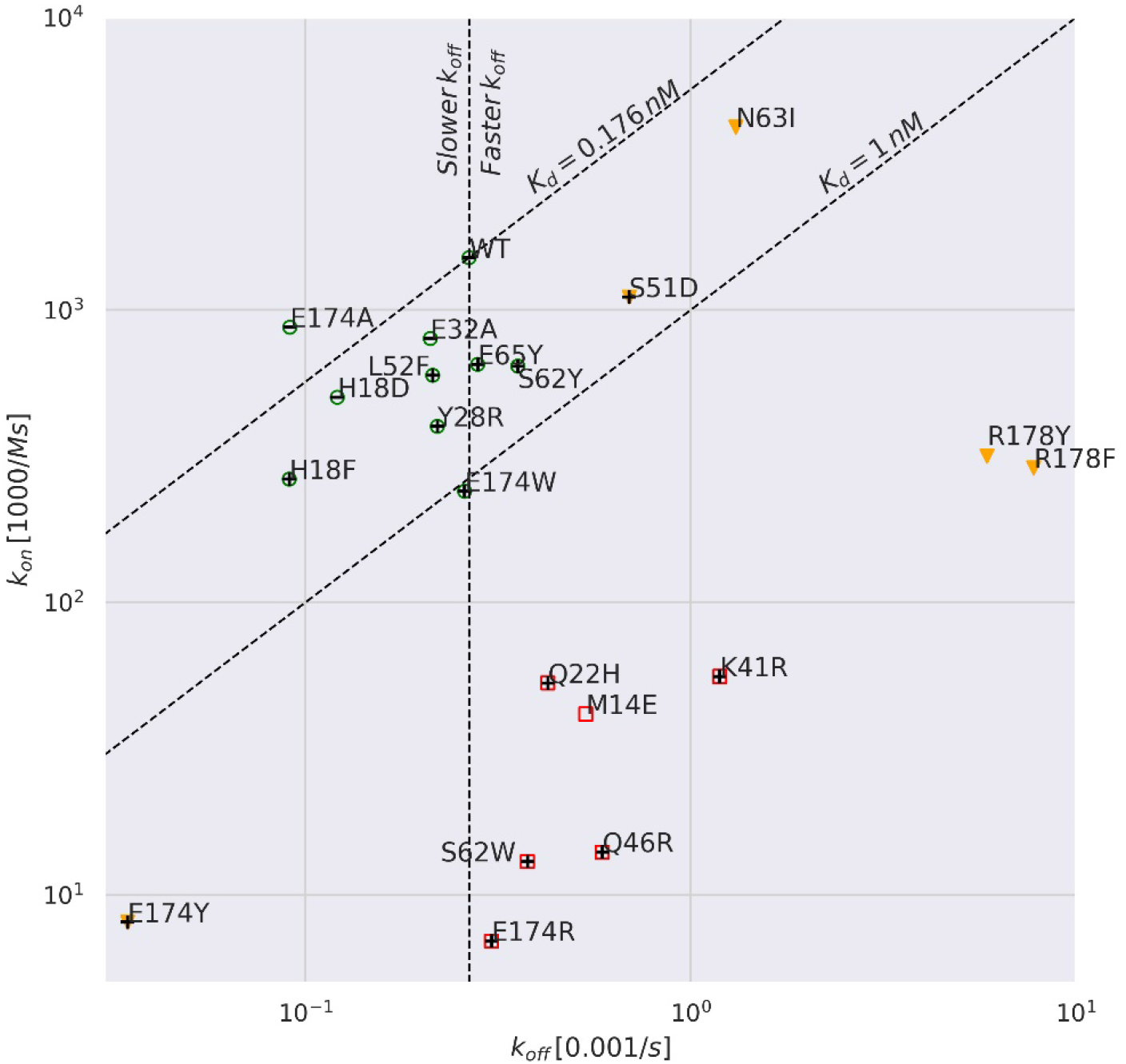
k_on_ vs k_off_.

### Surface Plasmon Resonance (SPR) measurements

The SPR measurements produced clear and reproducible k_on_ and k_off_. (given in **Error! Reference source not found**. and Table 1). The SPR sensograms are provided in Supplementary Figures S1-S3.

**Table 1.** Kinetics of hGH/hGHR binding, measured by Surface Plasmon Resonance, and ordered by k_off_. Green shading indicates k_off_ slower than that of WT. In these cases, k_on_ is also slower, such that the only E174A has K_D_ lower than that of WT.

| $k_{on}$ [1000/Ms] | $k_{off}$ [0.001/s] | $K_D$[nM] | Substitution |
| --- | --- | --- | --- |
| 600 | 1000000 | 2 | M14W |
| 300 | 8 | 30 | R178F |
| 300 | 6 | 20 | R178Y |
| 4000 | 1 | 0.3 | N63I |
| 60 | 1 | 20 | K41R |
| 1000 | 0.7 | 0.6 | S51D |
| 10 | 0.6 | 40 | Q46R |
| 40 | 0.5 | 10 | M14E |
| 50 | 0.4 | 8 | Q22H |
| 10 | 0.4 | 30 | S62W |
| 600 | 0.4 | 0.5 | S62Y |
| 7 | 0.3 | 40 | E174R |
| 700 | 0.3 | 0.4 | E65Y |
| 2000 | 0.3 | 0.2 | WT |
| 200 | 0.3 | 1 | E174W |
| 400 | 0.2 | 0.6 | Y28R |
| 600 | 0.2 | 0.4 | L52F |
| 800 | 0.2 | 0.3 | E32A |
| 500 | 0.1 | 0.3 | H18D |
| 900 | .09 | 0.1 | E174A |
| 300 | .09 | 0.3 | H18F |
| 8 | .03 | 4 | E174Y |

Our SPR experiments yielded an unexpected result. As mentioned, we produced 22 GH Variants: the wild-type (WT), positive control (E174A), and 20 novel mutants (including E174Y). In this text all comparisons of kinetic quantities (KD, k_on_, k_off_, etc.) are with respect to WT. Recall that the dissociation rate is:

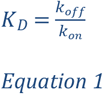

E174A had a much lower k_off_ but also a lower k_on_ -- the k_off_ won out leading to overall higher affinity (smaller K_d_) compared to WT. 13 of the novel mutants had significantly higher K_d_, and in particular k_off_ faster than WT. Six of the novel mutants (including E174Y) had slower k_off_ and k_on_, but unlike E174A in these six cases the slowdown in k_on_ won out and the end result was a higher K_d_. Interestingly, the highest-affinity mutants all decreased molecular mass (**Error! Reference source not found**.).

Dashed diagonals are iso-affinity contours (at WT and 1nM affinities). Vertical dashed line separates k_off_ slower on left (blue circles) from faster than WT on right (red squares). M14W is off the scale (very fast k_off_, low affinity). We identified a cluster of higher-affinity mutants (green circles) of which many (including E174A) have slower k_off_ than WT, but also slower k_on_. The slowest k_off_ belong to substitutions at zinc binding triad positions H18 and E174 (we did not mutate the third position, H21). We also identified a group with moderately faster k_off_ but much slower k_on_ (red squares). Mutants outside these two groups are marked with yellow triangles. “+” markers superimposed on the above indicate an increase in molecular mass, from WT to mutant, while “-” markers indicate a decrease in molecular mass. Substitutions with small mass changes have neither of these markers. Note that the three mass-decreasing mutants (E174A, H18D, E32A) are among the four highest-affinity mutants. N63I, the fourth mutant, also has a (slight) decrease in mass.

### Circular Dichroism (CD) measurements

The CD measurements feature a clear isodichroic point, for WT, L52F, and E174A/Y. (Figure 5) The salient difference between the four variants is the relative depths of the troughs at 208 and 222 nm, which may indicate a differential effect of E174Y and especially E174A on coiled-coil content. The differences in coiled-coiling reduce with temperature, and disappear around 40-45C.(Figure 6) L52F is not positioned to affect the coiled-coil structure and serves as a control for that purpose.

**Figure 5.**
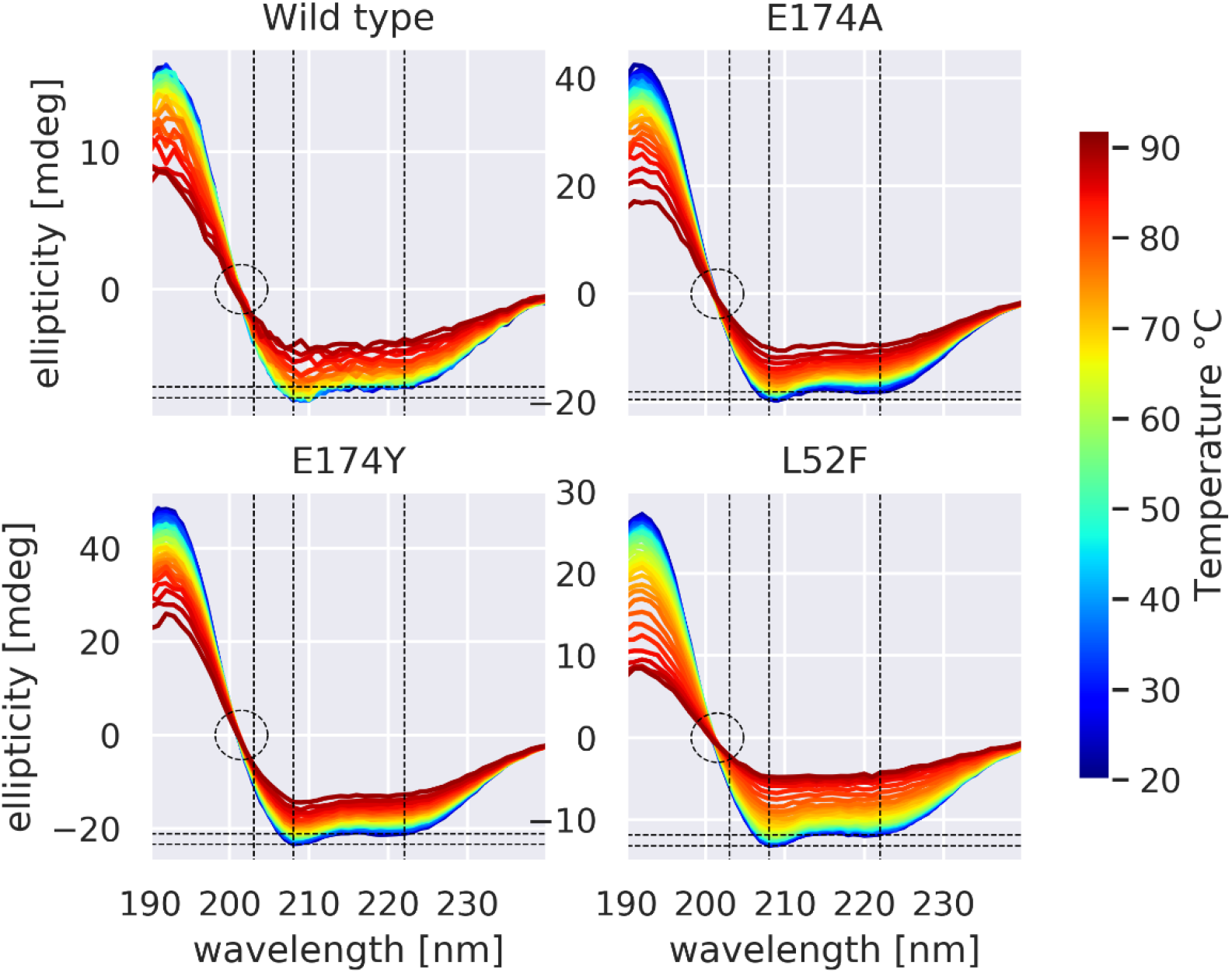

**Figure 6.**
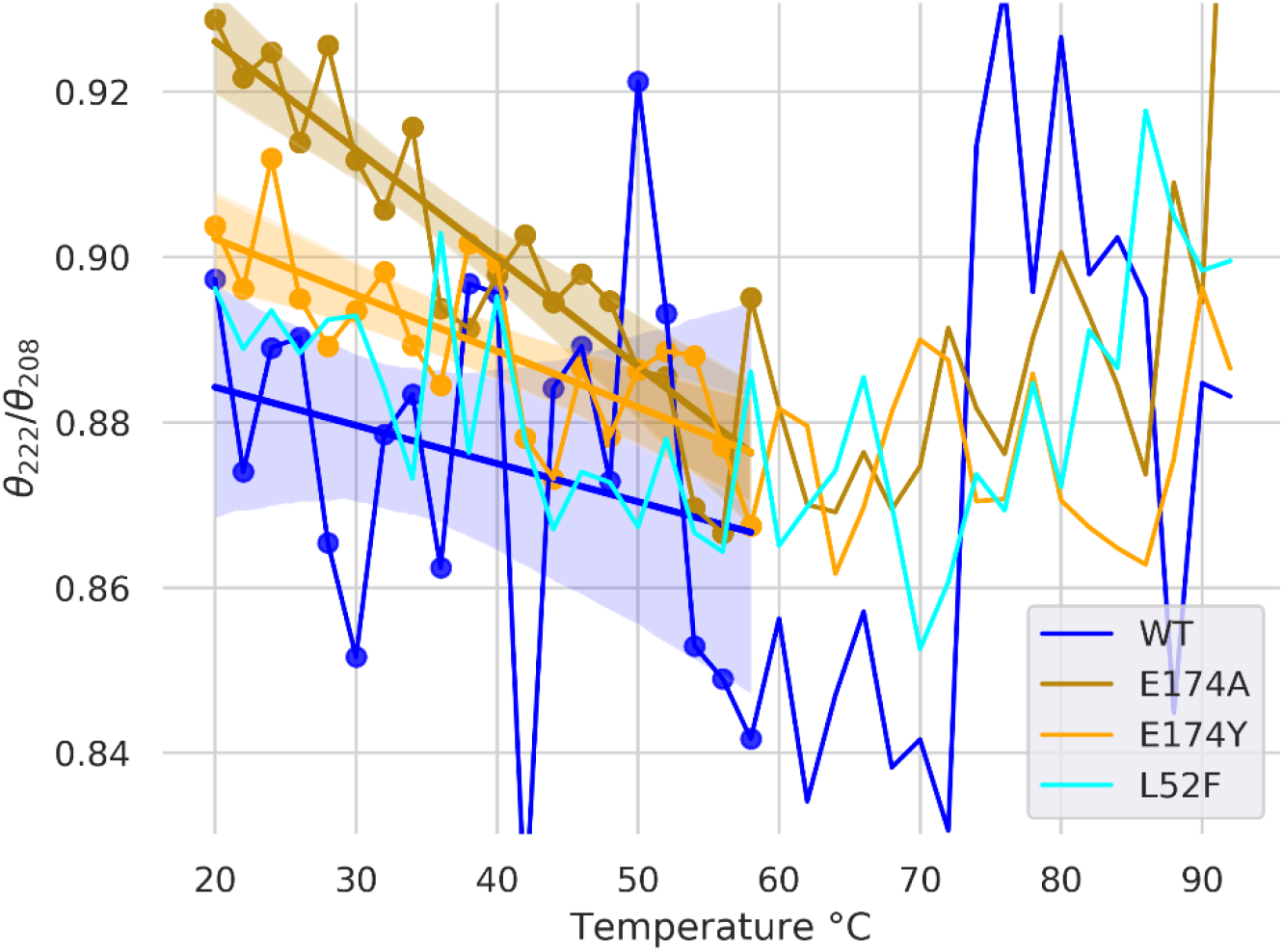

Circular Dichroism spectra for WT and three hGH variants, observed at temperatures from 20 to 92C. Note the presence of clear isodichroic points at 201.5 nm (circled), indicating only α-helical and random-coil content (no β sheets),(Scholtz *et al*., 1991) consistent with crystallographic data (note this is clearly below 203 nm). The maximum at 190 nm and minima at 208 and 222 nm likewise indicate alpha helices. Higher ratio of ellipticity at 222 vs 208 nm (θ_222_/θ_208_) is associated with higher coiled-coil content. Dashed vertical lines indicate 203, 209, and 222nm. Dashed horizontal lines indicate the ellipticity at these two wavelengths and 20C – the smaller relative gap for E174A (compared to WT, E174Y, and L52F) is visually apparent. The isodichroic point (zero crossing) indicates the protein is two-state, coil and helix (Holtzer and Holtzer, 1992); our isodichroic point is near the 201.5nm reported by (Correa and Ramos, 2009), rather than the 203-205nm reported elsewhere (Holtzer and Holtzer, 1992).

Mutations at position E174 affect θ_222_/θ_208_, suggesting increased coiled-coil content. We plotted θ_222_/θ_208_, for temperatures from 20 to 92C. Ratios greater than unity typically indicate coiled-coils. Θ_222_/θ_208_ did not exceed unity for E174A, which may be because only some of the helices are in a coiled-coil. θ_222_/θ_208_ lower than 0.9 indicates helices in isolation. (Crooks *et al*., 2011) Differences in ratio (comparing WT to E174A and E174Y) are greater at the lower end of the temperature range, presumably due to greater overall order. Linear regressions are shown for all variants (except for L52F, left out for clarity) over the temperature range of 20 to 58C (in which θ_222_/θ_208_ varies linearly). Also shown are 95% confidence intervals for the regressions (generated by the seaborn regplot function, in light shading). E174A has clearly higher θ_222_/θ_208_ than E174Y and WT. E174Y has somewhat higher θ_222_/θ_208_ than WT. L52F is a control, as it is not in a helix and not near the coiled-coil; its θ_222_/θ_208_ is indistinguishable from that of WT, and is less than that of E174A and possibly that of E174Y. E174A / E174Y may somewhat-increase degree of coiled-coiling compared to WT, while L52F has no effect.

## Discussion

The affinity increase induced by the E174A substitution (at 0.08nM, vs. 0.3nM for WT) was quite surprising at the time of its discovery (Cunningham and Wells, 1989) as alanine scanning usually disrupts rather than increasing binding. This position is in the zinc binding triad, and is highly conserved. In this work we confirm the earlier reported (Cunningham and Wells, 1989) increase in affinity of E174A, but now clarify that it is due to a decrease in k_off_ which more than counters a decrease in k_on_. The decrease in both quantities (albeit to varying degree), is shared by several other mutations. In the case of E174A/Y, the mutations may be increasing coiled-coilness, at least in the absence of zinc.

E174A better obeys the heptad pattern, compared to E174. The CD experiment was done with HPLC-purified hGH variants – so no ions were present. E174 is part of the zinc-binding triad a triad which by inspection would appear to stabilize the coiled-coil. So with physiological Zinc coiled-coilness may increase for E174. Zinc-induced hGH homodimerization also competes with hGH-hGHR binding, and E174A, by destroying zinc binding, may reduce the former to the benefit of the latter (when zinc is present). Note that our SPR experiments were done with hGH which was *not* HPLC-purified, and so presumably contain nickel from the nickel column, which could be bound by the triad. In the absence of zinc, E174A may increase coiled-coiling (e.g. by encouraging packing). Further experiments are needed to resolve this issue.

The results also speak to the dearth of good published results for affinity maturation by computational selection of substitutions at a single position, compared to the many successes of combinatorial techniques like phage display (Whitehead *et al*., 2013). As we have seen the former may change monomer properties, which the latter may compensate with substitutions at other positions. All potentials must balance the stabilizing effect of additional contacts, against the destabilizing effect of increased molecular volume. The latter can be due to entropy, desolvation, and sterics. The steric effect may be particularly important in our case since the mutations are mostly at alpha helical positions, which have little backbone flexibility. In (Flores *et al*., 2021), Dataset C, of 11 SKEMPI2 (Jankauskaite *et al*., 2019) mutations with ΔΔG_experimental_ < -0.7 kcal/mol, only three increased mass, and included two positions in coils (1KAC, 1JTG) and one in a very short helix on the interface rim (2GYK). Many mass-increasing mutations were selected by our method, while only the mass-decreasing mutations yielded near-WT affinities, thus FoldX appears not to get this balance right in the context of helices in the PPI core (Levy, 2010). In prior work we found near-WT affinities even with increases in molecular mass(Nosrati *et al*., 2017), but these were on the PPI rim (Levy, 2010), where there is more space available. Thus in future application of this method for that goal, it may be beneficial to explicitly select mass- or volume-decreasing substitutions for any positions in the core and on secondary-structural elements.

## Acknowledgements

We gratefully acknowledge support from the Swedish Foundation for International Cooperation in Research and Higher Education (STINT) and the Lars Hierta Memorial Foundation. S. Flores is funded by the Swedish Research Council grant 1024046, the National Research School in Medical Bioinformatics. The SciLifeLab Drug Discovery and Development Platform provided differential scanning fluorimetry equipment. Hugo Barrera provided biochemical and evolutionary advice. Faruck Morcos helped us look for a coevolutionary signal in some rather limited sequence alignments.

## References

Clackson, T., Ultsch, M.H., Wells, J. a and Vos, a M. de (1998) J. Mol. Biol., 277, 1111–28.

Correa, D.H.A. and Ramos, C.H.I. (2009) African J. Biochem. Res., 3, 164–173.

Crooks, R.O., Rao, T. and Mason, J.M. (2011) J. Biol. Chem.

Cunningham, B., Mulkerrin, M. and Wells, J. (1991a) Science (80-.)., 253, 545–548.

Cunningham, B. and Wells, J. (1989) Science (80-.)., 244, 1081–1085.

Cunningham, B.C., Ultsch, M., Vos, A.M. De, Mulkerrin, M.G., Clauser, K.R. and Wells, J.A. (1991b) Science (80-.)., 254, 821LP–825.

Danielson, U.H. (2009) http://dx.doi.org/10.4155/fmc.09.100, 1, 1399–1414.

Dourado, D.F.A.R. and Flores, S.C. (2014) Proteins, 82, 2681–90.

Flores, S.C., Alexiou, A. and Glaros, A. (2021) PLoS One, 16, e0257614.

Flyvbjerg, A., Bennett, W.F., Rasch, R., Kopchick, J.J. and Scarlett, J.A. (1999) Diabetes, 48, 377–82.

Guerois, R., Nielsen, J.E. and Serrano, L. (2002) J. Mol. Biol., 320, 369–87.

Holtzer, M.E. and Holtzer, A. (1992) Biopolymers, 32, 1675–1677.

Jankauskaite, J., Jiménez-García, B., Dapkunas, J., Fernández-Recio, J. and Moal, I.H. (2019) Bioinformatics, 35, 462–469.

Kim, M.J., Park, H.S., Seo, K.H., Yang, H.J., Kim, S.K. and Choi, J.H. (2013) PLoS One, 8, e56168.

Levy, E.D. (2010) J. Mol. Biol., 403, 660–70.

Lowman, H.B. and Wells, J.A. (1993) J. Mol. Biol., 234, 564–78.

Lupas, A., Dyke, M. Van and Stock, J. (1991) Science (80-.).

Nosrati, M., Solbak, S., Nordesjö, O., et al. (2017) Protein Eng. Des. Sel., 30.

Ross, R.J., Leung, K.C., Maamra, M., Bennett, W., Doyle, N., Waters, M.J. and Ho, K.K. (2001) J. Clin. Endocrinol. Metab., 86, 1716–23.

Scholtz, J.M., Qian, H., York, E.J., Stewart, J.M. and Baldwin, R.L. (1991) Biopolymers, 31, 1463–1470.

Whitehead, T.A., Baker, D. and Fleishman, S.J. (2013) Methods Enzymol., 523, 1–19.

